# Genome polarisation for detecting barriers to geneflow

**DOI:** 10.1101/2022.03.24.485605

**Authors:** Stuart J. E. Baird, Jan Petružela, Izar Jaroň, Pavel Škrabánek, Natália Martínková

## Abstract

1. Semi-permeable barriers to geneflow in principle allow distantly related organisms to capture and exchange pre-adapted genes potentially speeding adaptation. However, describing barriers to geneflow on a genomic scale is non-trivial.
2. We extend classic diagnostic allele counting measures of geneflow across a barrier to the case of genome-scale data. Diagnostic index expectation maximisation (*diem*) polarises the labelling of bistate markers with respect to the sides of a barrier. An initial state of ignorance is enforced by starting with randomly generated marker polarisations. This means there is no prior on population or taxon membership of the genomes concerned. Using a deterministic data labelling, small numbers of classic diagnostic markers can be replaced by large numbers of markers, each with a diagnostic index. Individuals’ hybrid indices (genome admixture proportions) are then calculated genome wide conditioned on marker diagnosticity; within diploid, haplodiploid and/or haploid genome compartments; or indeed over any subset of markers, allowing classical cline width/barrier strength comparisons along genomes. Along-genome barrier strength heterogeneity allows for barrier regions to be identified. Further, blocks of genetic material that have introgressed across a barrier are easily identified with high power.
3. *diem* indicates panmixis among *Myotis myotis* bat genomes, with a barrier separating low data quality outliers. In a *Mus musculus domesticus*/*Mus spretus* system, *diem* adds multiple introgressions of olfactory (and vomeronasal) gene clusters in one direction to previous demonstrations of a pesticide resistance gene introgressing in the opposite direction across a strong species barrier.
4. *diem* is a genomes analysis solution which scales over reduced representation genomics of thousands of markers to treatment of all variant sites in large genomes. While the method lends itself to visualisation, its output of markers with barrier-informative annotation will fuel research in population genetics, phylogenetics and association studies. *diem* can equip such downstream applications with millions of informative markers.

## Introduction

Divergence between taxa generates genome-wide correlations in SNP states (marker alleles). Geneflow across a barrier between taxa can be measured by counting alleles belonging on side A that have crossed to side B. The proportion of side A alleles an individual carries is its hybrid index; hybrid indices form clines across barriers from which we can estimate barrier strength (Barton 1979b,a). Classically, reference sets of ‘pure’ individuals from each side of a barrier were used to arbitrate where alleles belong: a marker fixed for different states in each reference set is diagnostic; which of its alleles belong on either side is considered proven. In modern terms, the use of diagnostic markers is reduced representation genomics favouring sites that enable geneflow to be detected. With genome scale sequencing we might hope to add many markers to a diagnostic set, but when adding a marker, a decision must be made regarding where its alleles belong.

While use of diagnostic markers has been described as introducing a bias against detection of geneflow (Gompert *et al*. 2017), this bias results only from declaring a priori that reference sets of individuals are ‘pure’. The critical decision regarding which side of a barrier an allele belongs does not require reference sets. For example, when Falush’s linkage model is used within the Structure inference framework (Falush *et al*. 2003a,b), the SITEBYSITE option produces a separate output file containing posterior population of origin assignments for each allele copy at each locus for each individual (Pritchard *et al*. 2010). This is not, however, a scalable solution, as convergence times under the linkage model increase rapidly with number of sites, even at *K* = 2. fineSTRUCTURE (Lawson *et al*. 2012) is more computationally efficient, but unlike Structure, requires declaration of ‘pure’ reference sets, and so re-introduces a bias against detection of geneflow.

Genome painting where alleles belong is difficult and computationally intensive using existing tools. Here we explore how classical reduced representation genomics for allele counting across a barrier can be adapted to high throughput sequencing datasets, and what measures characterising markers might be useful. We let correlation across the entire dataset arbitrate the allele/barrier-side decision. This is a data labelling task, and so open to expectation maximisation (EM) (Dempster *et al*. 1977).

## Materials and Methods

Consider a marker with two common states in a set of genomes: ‘A’ and ‘T’. If there is a barrier to geneflow within the genomes set, ‘A’ may be associated with genomes on one side, ‘T’ with those on the other side. Which way round these associations go is the *polarity* of the marker with respect to the barrier, and the evidential support for this decision can be calculated. We can represent an initial state of ignorance by using a random number generator to assign polarities to all markers. This is a *null polarisation*, and is equivalent to saying we work with no prior on population or taxon membership of the genomes. A null polarisation does not, however, remove information regarding associations between markers. If divergence has generated genome wide correlations in state, then a second marker with common states ‘C’ and ‘G’ will show ‘A’ genomes tend to have ‘C’ while ‘T’ genomes have ‘C’. Of course, the converse can be true. A null polarisation by definition will get about half these cross-locus associations correct. This is sufficient signal that some of the remaining markers can be given the matching polarisation. The diagnostic index expectation maximisation (*diem*) algorithm iteratively co-polarises remaining markers. This process accelerates as further signal of genome-wide association is pulled out of the data by an increasingly correct labelling of markers by barrier side. For a given initial null polarisation, *diem* will always halt at the same final marker labelling. These labelings can be used directly to visualise the nature of a barrier, and the labelled data is ideal for multiple downstream applications in population genomics.

The *diem* algorithm uses likelihood-based genome polarisation to find which alleles of genomic markers belong on which side of a barrier and as a result co-estimates which individuals belong on either side and which genome regions have introgressed (Fig. 1). A pseudocode description of the algorithm is provided in Algorithm 1.

**Figure 1:**
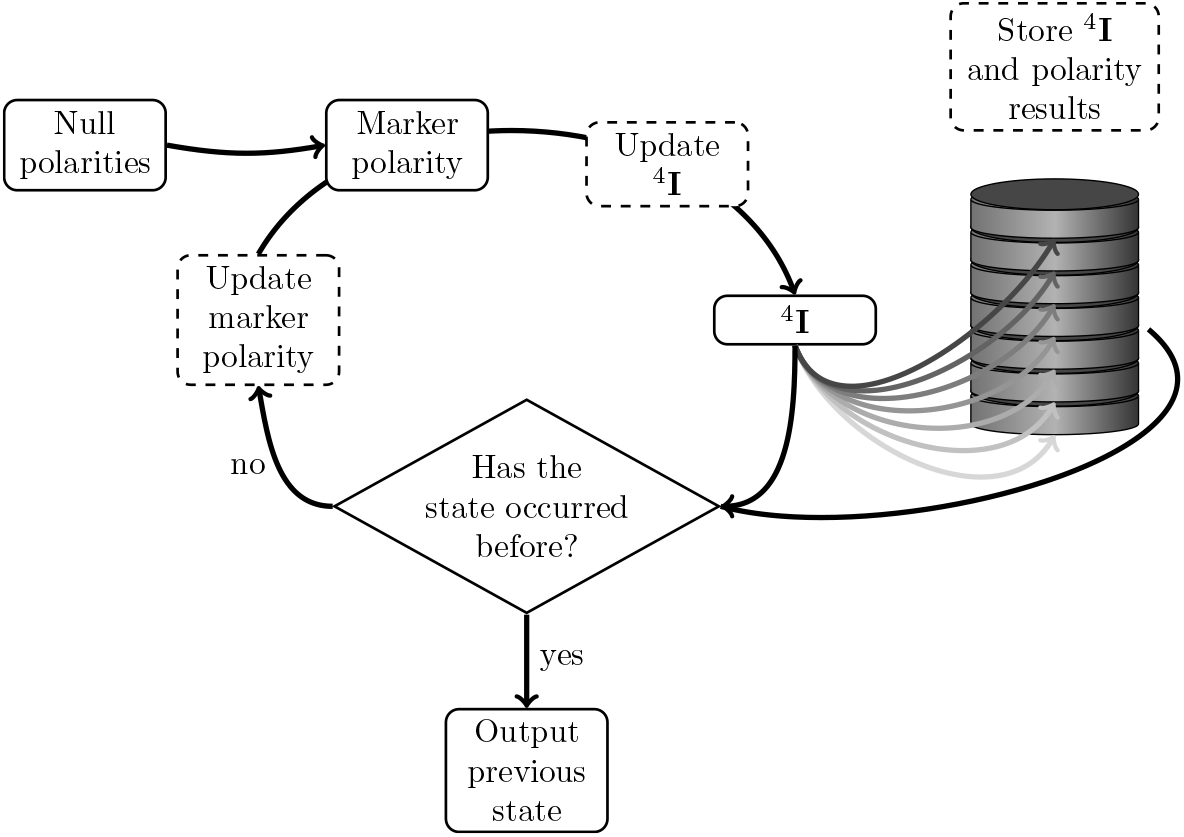
Diagnostic index expectation maximisation algorithm for detecting barriers to geneflow mplemented in diemmca, diempy, and diemr. ^4^**I** is a 4-genomic state count matrix (Eq. 4).

### The input data

At each site the two states which are most common across the genomes considered are identified. A toy dataset of three markers genotyped for six individuals has this form:

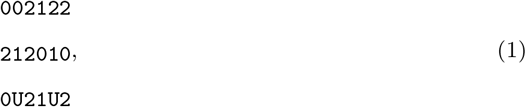

where genotypes encoded as 0 are homozygotes of one common state, 2 homozygotes of the other common state, 1 the common states heterozygotes, and U, genotypes which cannot be represented in terms of the two common states (naturally including the missing data case).

### Polarisation

For the first marker of the toy example (Eq. 1), suppose the two most common states are A and T. An alphabetic user encoding might encode homozygotes AA:0, TT:2. A major/minor allele user encoding might encode homozygotes AA:2, TT:0. The initial *diem* polarisation ‘flips’ (see below) half of user marker encodings at random, erasing any such encoding biases. The final *diem* polarisation flips a subset of encodings that maximises signed correlation of encoding across markers. There are always two mirror-image final polarisations. For the toy dataset, flipping first and final markers gives final polarisation

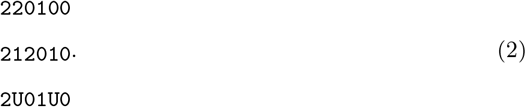

The mirror-image polarisation flips only the middle marker.

Within *diem*, a set of markers (in any original order) is then always encoded relative to the current polarisation as a matrix

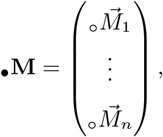

where *n* signifies the number of markers. 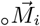 is the user encoded ith marker in the ordering of the marker set. The *diem* matrix •**M** will have dimensions n × *N*, where *n* is the number of markers and *N* is the number of individuals. For the toy example, the matrix •**M** is a 3 × 6 matrix.

For dataset •**M** of *n* markers, denote the current polarisation as 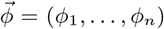, where *ϕ_i_* is the polarity of the ith marker, and *ϕ_i_ ∈* {0, 1} for ∀*i* ∈ {1, …, *n*}. The ith marker •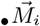 is encoded with polarity *ϕ_i_* as

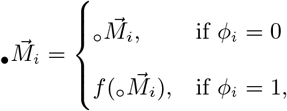

where the transition ◦ → • denotes the transition from user encoding to encoding with the current polarity *ϕ*. The function *f*(·) (‘flip’) flips the encoding of elements of the marker that correspond to the common state homozygotes. Genome states 0 and 2 change encoding to 2 and 0, respectively, while genome states 1 and U are unchanged.

The mirror-image (bitwise negation) final polarisations of the toy dataset are then 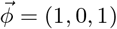 (Eq. 2), and 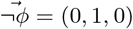 respectively.

### The genomic 4-state count matrix

We use a genome encoding that allows four states for each genotype of a marker in an individual. These genomic states are ordered U, 0, 1, and 2. Each marker 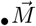 encoding can be expanded to a binary presence/absence matrix for the four genomic states in each individual. Individuals constitute rows, the genomic states constitute columns of the matrix. Each row sums to 1, because each individual has only one encoding for a marker. We will denote the transformation of the *i*th marker 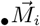 into the matrix 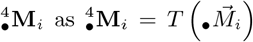. The first marker in the toy example (Eq. 1), under polarisation *ϕ*_1_ = 0, 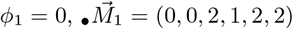 is transformed into the matrix

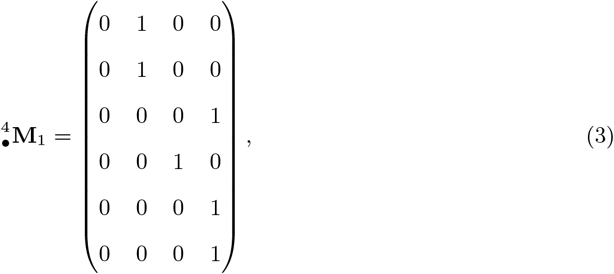

where the first column corresponds to the unknown state U, the second column to homozygote encoding 0, the third column to heterozygote encoding 1 and the fourth column to homozygote encoding 2.

When we sum the 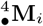 matrices across the whole genotyped dataset, we obtain an *N* × 4 matrix of genomic state counts

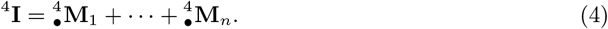

The rows in ^4^**I** refer to individuals and the elements of the matrix are integers in the range from 0 to *n*, each row summing to the number of markers *n*. This means that the whole genome of one individual is summarized as a single row of four values in ^4^**I**, and the entire genome set by ^4^**I**.

Hereon, the ^4^ prescript will refer to a vector or a matrix that represents the four states a genotype can have in *diem*. A matrix with the ^4^ prescript will have rows corresponding to individuals and columns to the genomic 4-state counts.

### Correcting for departures from large *n* markers assumptions

^4^**I** contains a summary of the whole dataset with which marker states across individuals may be correlated. For genome scale data, ^4^**I** could be used directly to calculate expected state frequencies, because in the limit of large *n* we can assume that first, all states will be observed for all individuals, and second, the contribution of any one marker to expectations is infinitessimal (i.e. negligible). To allow application over all data scales, we correct for departures from these large *n* assumptions. We will refer to this as ‘small data error correction’ *ζ*, as the correction depends on the smallest observed state count. With respect to the first assumption, if any individual has an unobserved state (count = 0), then no expectation for that state in that individual can be formed. As to the second assumption, if the number of markers *n* is small, then the contribution of any one marker to the expectation will no longer be negligible. These two departures define three cases of the small data error correction

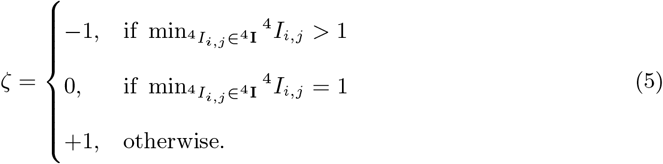

We add the small data error correction *ζ* to all elements of ^4^**I** to obtain a corrected genomic 4-state count matrix

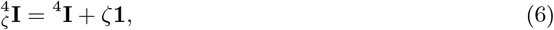

where **1** is a *N* × 4 all-ones matrix.

Small data error correction with *ζ* = +1 will be invoked for the toy dataset (Eq. 1) because, for example, not all individuals possess a U in their genomic states summary. Each individual’s genomic state summary is then augmented with one count of each possible state in 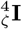. An analogy can be drawn with the Dirichlet prior on the multinomial distribution. The Dirichlet prior is flat when all state counts are 1. When ζ = +1, hybrid index, heterozygosity and error rate (all of which can be estimated from 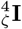, see below) must be interpreted with caution. *diem* outputs ζ in each iteration, allowing users to check for small data error correction effects in interpreting results. The ζ = +1 correction will introduce a bias in these indices. In genome scale data, invocation of this small data error correction will be rare and, if invoked, its effect will be small.

Where ζ = 0, the small data error correction leaves ^4^**I** unchanged. This case will be rare in genomic data, but is theoretically possible.

For most genome scale datasets, correction ζ will be equal to *−*1. By definition, more than one observed state of each marker is present, and we can efficiently remove the contribution of any marker in marker-expectation comparisons with ζ = *−*1 by inspection of Eqs. 3 and 4. Because 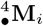 (Eq. 3) is always unit at the element positions of observed states, and otherwise zero, 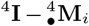 translates to 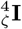 at the element positions of observed states. We will return to this computational efficiency later in the notation.

### Hybrid index, heterozygosity, error rate

Hybrid index *h*, observed heterozygosity *H*_O_ and error rate *E* can be estimated from 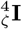. For a diploid genome compartment, the unconditional hybrid index for the ith individual is given as

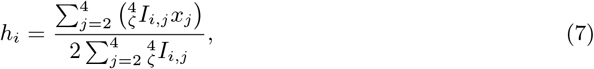

where *x_j_* is the *j*th element of a vector 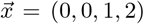 = (0, 0, 1, 2). The unconditional hybrid index of an individual is the proportion of alleles it carries that are associated with one side of the barrier when all markers are considered equally. Which side of the barrier depends on which of the mirror-image (bitwise negation) final polarisations *diem* converges to for a given initial polarisation. Only the last three columns in 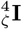 are used to calculate *h*, because the first column is undetermined genomic state count (U), and so cannot contribute to estimation of *h*.

The *diem* algorithm can be applied to diploid, haplodiploid and haploid marker sets. If marker data is known to come from genome compartments of different ploidy, *h* estimates should reflect that heterogeneity. When estimating *h*, the genomic 4-state count matrix ^4^**I** in Eq. 7 needs to be replaced by a genomic 4-state *allele* count matrix ^4^**A**. To calculate the ^4^**A** matrix, first a compartment-specific partial genomic 4-state count matrix is counted over each of the *k* compartments. For the *g*th compartment, the compartment-specific partial genomic 4-state count matrix is given as

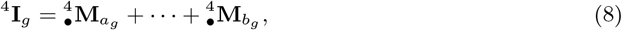

where *a_g_* and *b_g_* are indices specifying the positions of markers in the original marker ordering that describe the subset of markers belonging to the *g*th compartment.

The ^4^**A** matrix is a weighted sum of all *k* partial genomic 4-state count matrices with row values weighted by the ploidies of individuals of the specific compartment. For example male/females (m,f,m,f) for an X chromosome compartment have ploidies (1,2,1,2). The weighted sum is given as

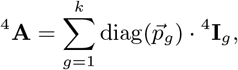

where 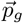 is the vector of individual ploidies for the *g*th compartment.

When ζ = +1, the small data correction needs to be distributed across the genomic compartments relative to the compartment sizes and ploidies.

The observed heterozygosity for the *i*th individual H_O_*i* is the proportion of heterozygous genotypes in the total of the called alleles in 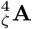 and is given as

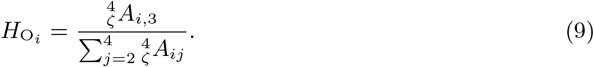

The error rate estimated from 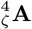 is the proportion of undetermined calls from the total number of genotype calls. For the *i*th individual, the error rate is

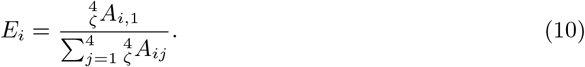

The observed heterozygosity and the error rate are independent of the data polarisation. An initial (random) null polarisation is expected to give *h* ≈ 0.5 for all genomes. Final hybrid index estimates are expected to be heterogeneous 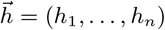 in the presence of a barrier, and will ‘flip’ 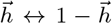 depending on which of the mirror-image (bitwise negation) final polarisations *diem* converges to for a given initial polarisation.

### Model of an ideal diagnostic marker

When considering the Platonic ideals of ‘pure’ individuals and (perfectly) ‘diagnostic’ markers, two operations applied to the hybrid index become useful: rescaling and ordering. This is because ‘pure’ individuals will have one of two hybrid indices: *h* ∈ {0, 1}, and a perfectly diagnostic marker will switch from one state to the other at the steepest change in hybrid index. We use unity-based normalization to calculate rescaled hybrid indexes *h*′. For the ith individual, the rescaled hybrid index is given as

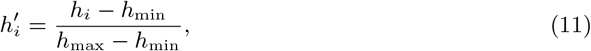

where *h*_max_ and *h*_min_ are the highest and the lowest values in 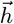, respectively. The 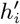 values are then ordered to form 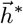 so that 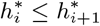.

To determine changes in the hybrid index, we calculate differences between neighbouring values of the rescaled and ordered hybrid indices in 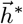. The ith element of a vector of the differences 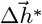 is given as

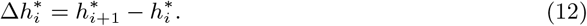

We identify the position of the largest difference in 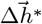 according to

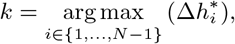

We estimate the barrier as lying in that interval between the (hybrid index ordered) individuals that shows this largest change in hybrid index. Then, a value of the hybrid index that splits individuals by barrier side can be calculated by interpolation of *h*^*′*^ as

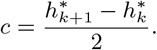

We can now construct an ideal diagnostic marker 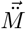, where the *i*th individual will have the ideal genotype

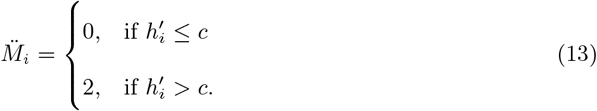

We can now construct an ideal genomic 4-state count matrix 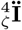 that would arise if all markers were perfectly diagnostic. To construct 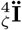, we first need to transform the vector 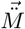 into a 4-genomic state matrix 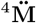 (cf Eq. 3). The ideal 4-genomic state count matrix 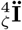 is then

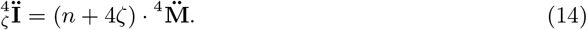

On the metric of rescaled hybrid indices *h*^*′*^, each individual lies at some positive or negative displacement from *c*. We define a weight *β* ∈ (0, 1] which increases linearly with the magnitude of this displacement. For the *i*th individual, the weight is given as

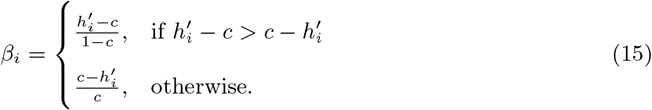

Individuals with extreme hybrid indices, i.e. individuals with *h* ∈ {0, 1} (tending to Platonic ‘purity’), have high weight, those with intermediate hybrid indices have low weight. Under either final polarisation of the toy example, the first and last individuals will have maximal weight *β* = 1. We use these weights to construct a linear combination ^4^**V** of the observed genomic 4-state counts 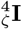 and the ideal genomic 4-state count matrix 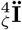 as

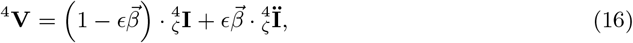

where *ϵ* is an error term determining how much we want the ideal ideal genomic 4-state count 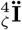 to influence the analysis, or conversely how much we wish the nature of real data to impinge on the ideal model.

### Likelihood of a marker’s polarity encoded state

The log-likelihood log *L* of observing a marker 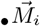 in a genomic dataset is calculated conditional on the frequencies of the four states in each individual across all other markers. A matrix of these genomic state frequencies is given as

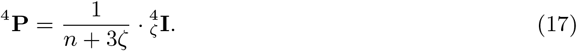

The log L of the ith marker 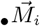 on 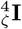 is then given as

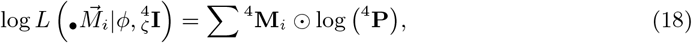

where ⊙ stands for the Hadamard product.

The base of the logarithm is arbitrary, and we will use a natural logarithm hereon. The log-likelihood will thus be ln *L*.

To this point, the likelihood calculation is model free; it is based only on the frequencies of observed states. For a set of genomes with majority U state, the most likely state will be U. The model of an ideal marker (and so the weighting toward ‘pure’ individuals) is introduced by substituting ^4^**V** for 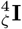 in the calculation of 4-genomic state frequencies. These become frequencies expected given the model, and the likelihood calculation is given as

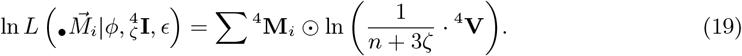

Now, for *ϵ* = 0, the likelihood is model free. For 0 < *ϵ* < 1, the likelihood is influenced by the model and at *ϵ* = 1, the likelihood is for the Platonic ideal model that does not admit U or 1 genomic states. We leave ϵ as a free parameter, but recommend it is set close to 1. Then U,1 states are permissible, with likelihoods proportional to their frequency of occurrence in the data.

### Diagnostic index, support and polarity

The outputs of *diem* are diagnostic indices *d*, supports *δ*, and polarities *ϕ* for each marker. At initialisation, the user chooses the error term *ϵ* (0.99999 is recommended), and the initial polarisation of all markers 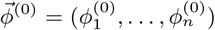 (a null of random initial polarities is recommended). The algorithm proceeds by iterative expectation maximisation (EM). At each iteration, the current polarity *ϕ* is stored. Likelihoods (Eq. 19) of the two alternative polarity encodings for each marker are compared. The maximum likelihood (ML) polarity is identified given the expected encoding state frequencies from counts across all other markers, as parameterised by *ϵ*.

In the jth iteration, the diagnostic index and support for the ith marker are given as

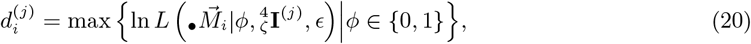

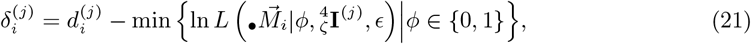

where 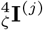 is the corrected 4-genomic state count matrix for the (*j* − 1)th polarity of the ith marker 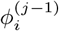.

The resulting polarity of the *i*th marker from the *j*th EM iteration is given as

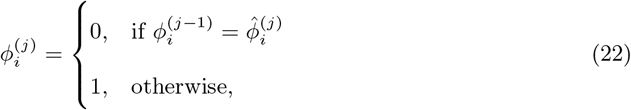

where 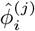 is the ML marker polarity for the *j*th iteration according to

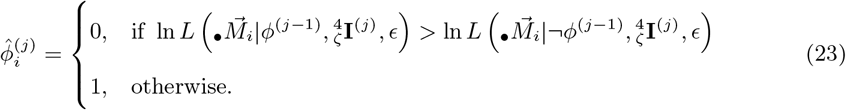

It is important to note that the calculations for each marker are independent (can be carried out in any order), allowing the computation for each iteration to be parallelised. The algorithm halts when the current polarity, or its bitwise negation, are found to exist in the store of polarities. That is, the EM halts in the jth iteration if *∃l* ∈ {0, …, *j* − 1} such that 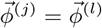. *diem* 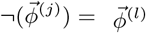 *diem* has then either converged or entered a limit cycle. Diagnostic index, support and polarity are then output for each marker with the same ordering as for marker input.

#### Algorithm 1 The *diem* algorithm in the case of homogeneous ploidy.

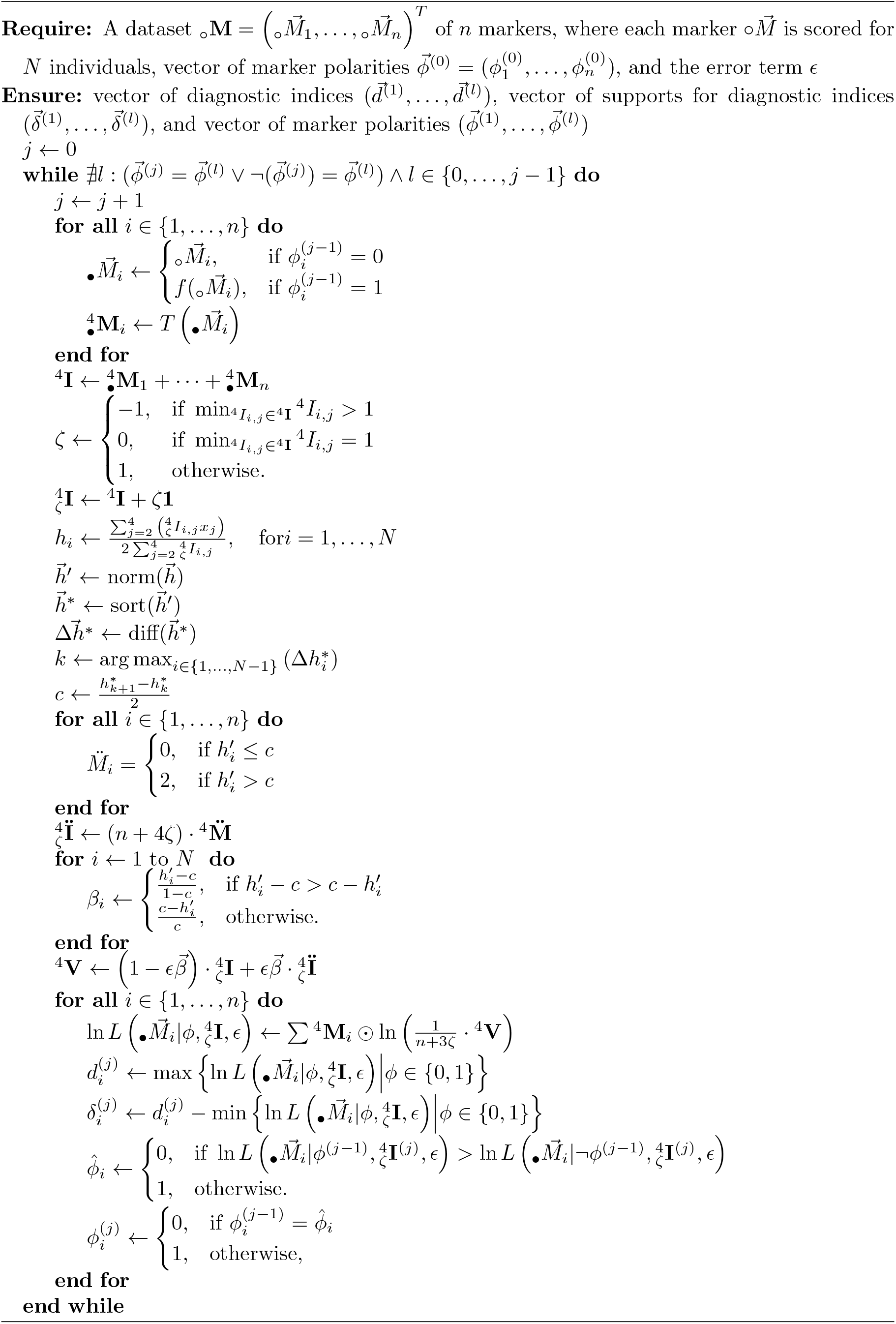

### Performance optimisation of EM iterations

In each EM iteration, all data are encoded according to the current polarisation *ϕ*. To do so, *diem* reads user encoded data (see section Polarisation) from store. For large data, this ‘read from store’ operation can be costly. *diem* addresses this issue in two ways. First, *diem* uses a compact data storage, where user encoded data for each marker is encoded as a string “012U21…”. Second, *diem* uses parallel data input, where user encoded data compartments are read in parallel (multi-core processors have multiple buffered pipelines to/from storage).

In each EM iteration, the 4-genomic state count matrix ^4^**I** must be calculated. In the first EM iteration there is no way to do so other than counting all encoded states. Thereafter, the same outcome is made more efficient by ‘Delta updating’ (cf. Delta encoding). To update ^4^**I** with respect to the current polarisation, *diem* first calculates a partial 4-genomic state matrix 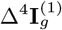 for each compartment. The equation is analogous to Eq. 4, but only markers with changed polarity are included, i.e.

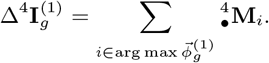

This partial count is sufficient information to update the second and fourth columns of ^4^**I** because 0, 2 encoding changes ‘flip’ counts between only these columns. The updating procedure is as follows:

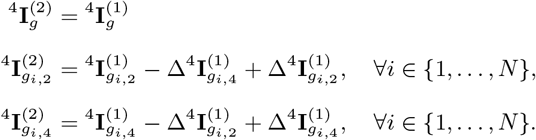

The *diem* likelihood calculation is required 2×n times for each iteration (cf. Eq. 23). However, the non-zero positions in 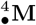 of a marker are invariant and need be recorded only once during *diem* initialization. Flattening the values ln 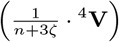 to a vector, the likelihood calculation for a marker (Eq. 19) reduces to a non-zero position lookup of that vector, which is then summed. Note that the frequencies of the weighted 4-genomic state count matrix (^4^**V**) are calculated by adding 3ζ to the denominator (Eq. 19). The operation removes the contribution of the marker *i* from the likelihood calculation to evaluate the likelihood of observing the state of the marker given the states of all *other* markers.

### Performance optimisation for detection of the halting state

The EM algorithm halts when the current polarity state (or its bitwise negation dual) has been visited before. For large data, this condition requires comparison of the current polarity with increasingly many stored polarities. Each is a comparison between two boolean vectors with length *n*, the number of markers. For high *n* and iterations *j* this brings a computational cost that can be reduced without loss of accuracy.

We assert that it is not possible for 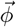 to satisfy the halting condition without ^4^**I** also recurring in value during EM iterations. Comparing current ^4^**I** with a stored ^4^**I** requires 4N integer comparisons, which tends to be more compact than *n* comparisons of the current and stored 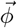 for genome-wide data. Therefore, when testing for the halting condition, we first test for recurrence of ^4^**I** (or its flipped mirror image, cf. end of section Hybrid index, heterozygosity, error rate). The cost of the full test for polarity recurrence in 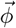 is only imposed if ^4^**I** recurrence is already shown.

## Alternative perspectives on the algorithm

This cross-disciplinary collaboration required consensus on a singular notation with which to describe *diem*. Debate centered on algorithmic versus analytic perspectives and conventions. In this section we briefly describe alternative perspectives that may be obscured by our notational choices, in the hope this will increase communication of the underlying science.

### *diem* as a Markovian finite state automaton

From an algorithmic point of view, *diem* is proposed as a closed form algorithmic solution: a halting finite state automaton, state 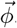. The number of distinguishable states is 2*^n^* because 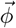 is binary, length *n*. Distinguishable states can be thought of as arranged along the finite number line from 1 to 2*^n^*. These states are however in equivalence duals with their bitwise negations: reordering states, we can think of the number line as having a fold at its centre. We can therefore label the finite automaton with a current state *S* in denary (decimal) *S* ∈ {1, …, 2^n*−*1^}. The toy dataset states can be labelled thus 1,2,3,4. The automaton state sampling is Markovian because it uses no memory of previous states. A Markovian random walk on a finite number line is guaranteed to revisit its start state, the *diem* halting requirement. A Markovian random walk on a folded finite number will do so more quickly. But the *diem* finite automaton does not carry out a random walk, its path is deterministic and guided by (conditioned on) a constant: the user encoded data. It is reassuring, however, that if all attempts at such conditioning fail, *diem* will still halt. It might be thought that *diem* is not Markovian because it holds the set of all states it has visited. This is incorrect, because that set is only referred to for the halting decision, never any state transition decision.

### *diem* as a hill climbing algorithm

*diem* is a strict hill climbing likelihood maximisation algorithm, but the grainsize of the strict hill climbing is one marker, so the number of axes of movement is *n* markers. As Fisher argued with regard to adaptive landscapes, *“In n dimensions only about one in* 2^n^ *[points can] be expected to be surrounded by lower ground in all directions”* (Fisher’s letter to Ford; Fisher *et al*. 1983, p. 201–202). The probability of a hill climber becoming trapped at a local optimum would then be low for large *n*, but this not a proof that *diem* will always find a global optimum. Such a proof depends not on the nature of the state space of the automaton, which is data dependent, but the rules that determine state transitions. These rules are formalised in the likelihood inference framework governed by data probabilities given expectations of the data. We judge these rules are best formalised with analytic rather than algorithmic notation, as the operation of constructing an expectation is one of modelling, and the gold standard of modelling is closed form analytic solution. We have therefore largely followed analytic notational conventions up to the current section, while trying also to place flags where notational conventions might differ. We will now revisit these flags while viewing the algorithm from further differing perspectives.

### *diem* as an operation on a set: No marker genome positions required

The most obvious strain between analytic and algorithmic conventions for *diem* is indexing. When an algorithm is applied over a set, the algorithmic notational convention is not to use indexing, as this implies an ordering on a set. If set members are exchangeable with respect to the algorithm, then it suffices to prove for any member. *diem* is applied to a set of markers, and no *diem* operation assumes they have any ordering. The memberships of markers in compartmental subsets again implies no ordering. The distinction of compartments by ploidy vectors over individuals again implies no ordering. Why then do we attach indices to markers in our notation? The answer is subtle. To operate, *diem* requires some (arbitrary, user chosen) orderings on both markers and individuals. These choices make no difference to the outcome. Our notation’s indexing of markers corresponds to one user-chosen ordering. Where markers can be ordered with respect to estimated positions in some reference genome, that would provide a natural ordering choice for users. We could have opened our methods description by explaining indices 1, …, *n* are best thought of as references to some arbitrary ordering. We chose not to.

### *diem* as an operation over ploidy compartments

The above diplomatically buried aspect of set operation notation resurfaces in Eq. 8. A natural indexed notation for one compartment of the markers is a subrange of their indices. This only makes sense however if we have chosen a non-arbitrary ordering of the markers. There are good algorithmic efficiency arguments for ordering makers such that those of similar ploidy lie in contiguous indexing subranges. There are also good algorithmic notational arguments for once again not referring to markers within these ploidy-equivalence sets by indices. Most obviously: an operation *m* can be parallelised with low overheads over ploidy equivanences *P*, marker memberships *S* and this parallelisation is implicit in the notation m(*P; S*). To express such an absence of an ordering requirement in our indexed notation, we contrast a **for** loop over (small) *n* where parallelisation would have little benefit with **for all** loops over (potentially very large) *n* where parallelisation benefits are clear (Algorithm 1).

### *diem* as a diagnostic labelling algorithm

Users may observe that *diem* replicates run on the same data but from different starting points (random nulls) give not one, but two answers. Why? *diem* operates without prior labelling of individuals (genomes). All meta data (taxonomic, geographic, phenotypic) is stripped off, leaving only ploidy patterns and encoded genetic state as inputs to the algorithm. And yet, if there is signal of a barrier in the genome data, *diem* will output an ordering on the individuals: a hybrid index. This one item is *both* of the observed *diem* answers. It is a Schrödinger’s hybrid index because it both runs from 0 to 1 and runs from 1 to 0 until the moment when it is observed re-united with the meta data. Then, from east to west, from taxon A to taxon B, from valley bottom to mountain top, the hybrid index runs either up, or down, *not* both. This is the limit of Gompert et al.’s (2017) assertion that diagnostic markers bias against detection of introgression. Diagnosis using imperfect prior information on what individuals are ‘pure’, and how to define ‘pure’ leads to a bias against identifying introgression. An absence of prior information has no imperfections, and cannot lead to such a bias. Diagnosis in the absence of prior information is unbiased, and the reduction in what we can infer working without prior is one bit: the up/down state of Schrödinger’s hybrid index.

### Implementation of the *diem* algorithm

The *diem* algorithm was developed in Mathematica as diemmca and idiomatically translated to Python, forming package diempy, and to R, forming package diemr. The diemmca and diempy implement *diem* genome polarisation including calculating hybrid indices, heterozygosity, genotyping error rate, the model of an ideal diagnostic marker (^4^**V**), and running the *diem* iterations to determine optimal marker polarities. The diemr package also implements a check on input data format, and polarized genomes can be exported/imported and visualized.

### Parallelisation grain

For UNIX-based operating systems, all *diem* implementations spread the computational load of processing genomic compartments (such as marker sets on chromosomes or scaffolds) over multiple cores. When the last core is finished, the core reports are summarized for genome-wide updates. This is in addition to the aforementioned parallelisation of read-from-store operations. All parallel operations have the same user-chosen *grain*: the compartments into which the set of input markers are divided.

## Validation

We cross-validated *diem* implementation in diemr and diemmca on two datasets. The first dataset consisted of 115 *Myotis myotis* bats scored for 31,074 SNP markers (Harazim *et al*. 2021, raw reads available at NCBI BioProject Accession: PRJNA681157). The bat data was chosen because current taxonomy, phylogeography and behavioural observation suggest there should be no barrier within this set of genomes, while the size of the dataset is representative of standard reduced representation genomics.

The second dataset included transcriptomic and genomic reads of 69 mice (36 *Mus musculus domesticus* and 33 *Mus spretus*, Supplementary Table S1) mapped onto mouse reference GRCm39, NCBI Assembly Accession: GCA 000001635.9. The full mouse dataset was of 38 million variable sites, and we used 117,403 SNPs at chr7:123189278–131234739 as a focus for comparative analyses. The *diem* implementation in diempy was cross validated with diemmca for the mouse dataset. The mouse system on which we demonstrate *diem* was chosen for a number of reasons. First, Sanger sequencing has shown that the worldwide human commensal *M. musculus domesticus* has captured a warfarin (pesticide) resistance gene, *Vkorc1*, from a non-commensal mouse species with ∼ 5% divergence, *Mus spretus* (Orth *et al*. 2002; Song *et al*. 2011). Second, genomic and transcriptomic high throughput sequencing (HTS) data have been made publicly available for this system for a decade. Third, the number of *domesticus/spretus* variable sites (38 million) exceeds the number of sites expected to vary between humans and Neanderthals (*∼*15 million). Demonstrating that *diem* scales to this mouse system is proof the method will scale to popular issues of hominin genome admixture. Fourth, the *domesticus/spretus* HTS data was recently analysed by co-authors of the original Sanger sequencing work (Banker *et al*. 2022).

## Comparison with existing methods

We compare *diem* to two methods inspired by phylogenetics (Banker *et al*. 2022) and population genetics (Falush *et al*. 2003a,b). We used partial mouse chromosome 7 region that is known to include a several-Mbp-long *spretus* introgression into a *domesticus* genome (Song *et al*. 2011).

Banker *et al*. (2022) used a sliding window sequence divergence-metric analysis to detect *domesticus*/*spretus* introgression for a subset of the ENA accessions we examine here. While we align genomes under a single reference (GRCm39, accession: GCA 000001635.9, assembled using long reads), the Banker *et al*. (2022) study involves distances measured relative to two different references: GRCm38 (aka mm10, short reads, proxy for *domesticus*) and GCA 001624865.1 (short reads, proxy for *spretus*). Direct comparison of the approaches’ output is made possible by NCBI Genome Remapping Service (https://www.ncbi.nlm.nih.gov/genome/tools/remap), which allows us to exchange Banker *et al*. (2022) introgression locations in their Supplementary Table 3 very precisely between reference accessions.

The second method was the Structure linkage model with the SITEBYSITE option (Falush *et al*. 2003a,b; Pritchard *et al*. 2010) that clusters individuals while allowing for admixture or hybridization between linked markers. We ran Structure for 10,000 generations following a 5,000-generation burnin.

## Results

We demonstrate *diem* on publicly available datasets: 31,074 SNPs from 115 bat genomes, and 38 million markers in 69 individuals for a genome barrier between species of mice.

### Bat SNPs dataset

Polarisation made little visual labelling difference in comparison to null in the bats dataset (Fig. 2A vs B). Pre-polarisation, hybrid indices were uniform *≈* 0.5 (Fig. 2C) as expected for a random null. Post-polarisation, hybrid indices run from 0 to ≈ 0.2 (Fig. 2C). Having found optimal marker polarities using *diem*, we conclude there is little or no evidence for a barrier to geneflow. We include a plot of diagnostic indices and their polarisation supports here (Fig. 2D) to demonstrate that, even in the case of no barrier, supports (statistically comparable across markers) correlate strongly with diagnostic indices (technically not comparable). This ‘shark’s fin’ shape is typical across all current analyses.

**Figure 2:**
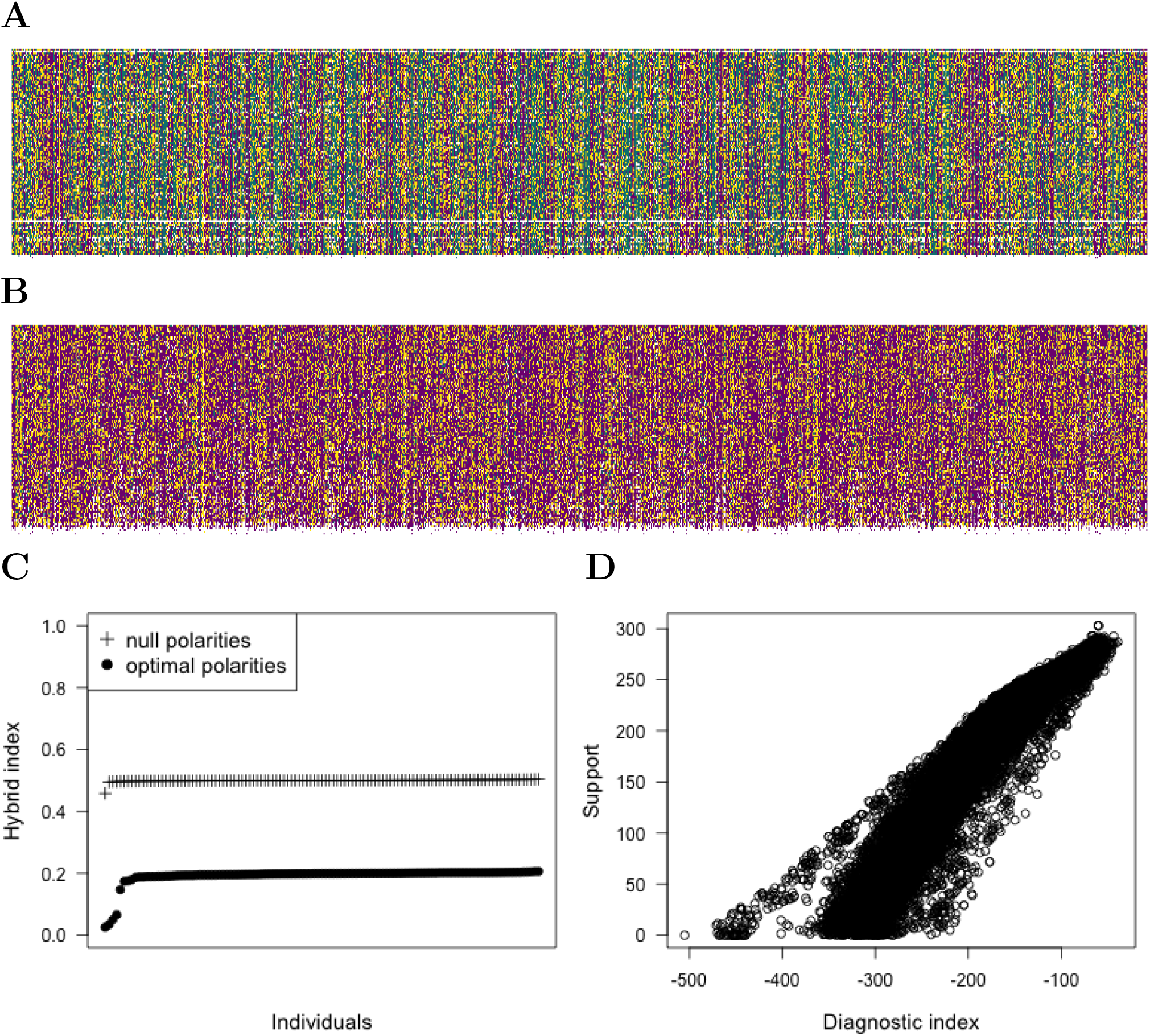
Genotypes of 31,074 SNP markers (columns) in 115 *Myotis myotis* bats (rows) (Harazim *et al*. 2021) with random null polarities (A), and with optimal polarities (B). Individuals in panel B are ordered according to their hybrid index. Purple – homozygotes for the allele attributed to barrier side A, green – homozygotes for the allele attributed to barrier side B, yellow – heterozygotes, white – unknown genotype state. C) Ordered individual hybrid indices for genotypes displayed in panels A and B. The optimal marker polarisation (circles) shows that the dataset consists of genetically similar individuals with the exception of five individuals with large proportion of unknown genotypes. D) Given a genome polarisation, the diagnostic indices of markers are strongly correlated with the supports for those markers’ polarisations. This is despite the fact that the former, being likelihoods, are therefore not strictly comparable across markers, while the later, being likelihood comparisons, are strictly comparable.

These results are consistent with the current taxonomy of *M. myotis*, and despite geographically widespread sampling. Identifying a single set of individuals is an important feature of *diem*. Unlike Structure (Pritchard *et al*. 2010), the *K* = 1 (no barrier) and *K* = 2 (barrier) cases can be distinguished by *diem*. The *diem* labelling of the bat dataset consists of a homogeneous set of genomes with five outliers. The outliers have relatively large numbers of unknown marker states (Fig. 2B). These states do not affect *diem* polarisation, rather, they co-occur with a shared homozygous marker pattern.

### Mouse whole-genome dataset

The visualization of the mouse genomes highlights the difference between individuals sequenced for whole genomes (horizontal blocks) and for transcriptomes (sparse data in vertical blocks above and below the whole-genome sequences) (Fig. 3A). The polarized genomic data (Fig. 3B) differentiate individuals with respect to the barrier to geneflow resulting in hybrid indices between

**Figure 3:**
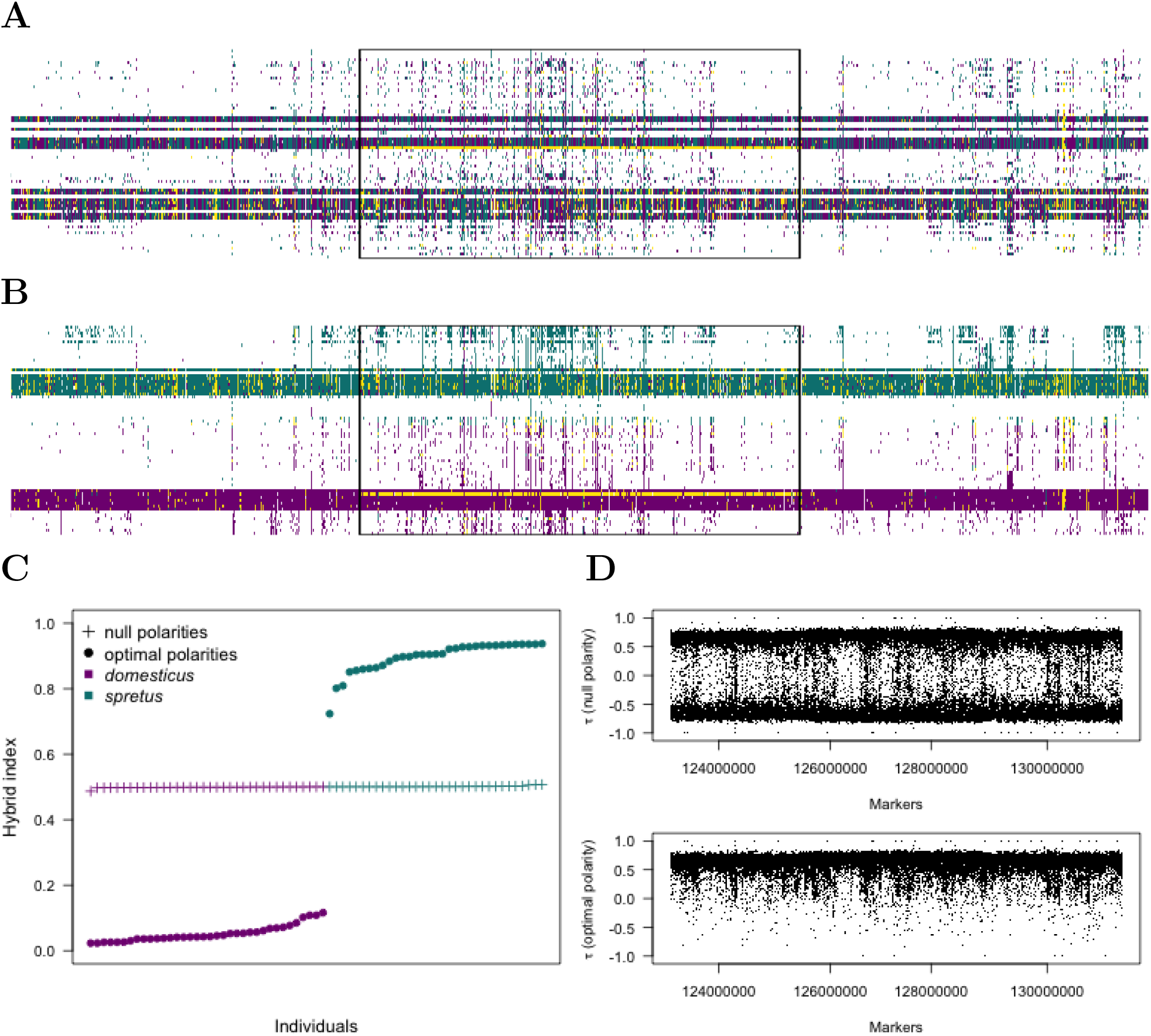
Genotypes of 117,403 markers (columns) on chromosome 7, positions 123,189,278–131,234,739 in 69 mice (rows) with random null polarities (A), and with optimal polarities (B). ndividuals in panel B are ordered according to their genome-wide hybrid index. The rectangle delimitates the genomic block carrying warfarin resistance that includes *Vkorc1* introgressed from *M. spretus* to *M. musculus domesticus* (Song *et al*. 2011). Two individuals have the introgressed block in our dataset. One is predominantly heterozygous, and the other is homozygous. Purple – homozygotes for the allele attributed to barrier side A (*domesticus*), green – homozygotes for the allele attributed to barrier side B (*spretus*), yellow – heterozygotes, white – unknown genotype state. Ordered individual hybrid indices for whole-genome markers in mice. The optimal marker polarisation (circles) shows that the dataset consists of two main groups of genetically similar individuals. Kendall’s τ correlation between hybrid indices and marker states in the genomic region displayed *n* panels A and B. The correlation coefficient is bimodal for markers with null polarity (upper panel) and unimodal for markers with optimal polarity with respect to barrier to geneflow (lower panel). ntrogression in two individuals is not apparent in this genome region scan.

0.02 and 0.93 (Fig. 3C). The Linnean binomial taxon labels separate at the steepest change in the hybrid index. In Fig. 3B, we show the introgressed region, where two *M. musculus domesticus* individuals carry the introgressed block previously identified by Sanger sequencing (Song *et al*. 2011). The introgression present in two individuals cannot be distinguished in a genomic scan correlating marker state to the individual hybrid index (Fig. 3D), but is obvious in the visualization of the polarised genomes (Fig. 3B).

### Implementation validation

Using identical initial null polarities, we found 0 differences in converged *diem* marker polarities in the bats and mice datasets as implemented in R, Mathematica, and Python. As the *diem* algorithm is deterministic, alternate implementations should give the same output on the same input. The diemr vs diemmca, and diempy vs diemmca, implementations were cross-validated in this way on multiple datasets, including large. Finding no output differences, the chance of coding error in any implementation is then the chance that all have coding errors with perfectly matched effect despite idiomatic translation into different languages. While the algorithm is concise, it is of sufficient complexity that accidental perfect error matching of this sort seems far fetched.

### Benchmarking

We benchmarked the *diem* algorithm implemented in diemmca on a MacBook Pro (Retina, 15-inch, Mid 2015), processor 2.8 GHz Quad-Core Intel Core i7, memory 16 GB 1600 MHz DDR3, 1Tb flash storage and diempy on a computational cluster (MetaCentrum). We used the full mouse dataset (all reference scaffolds), thinned at random to 10 reduced levels and ran *diem* for up to ten null initial polarisations for each level. The results show that the algorithm scales linearly with *n* markers (Fig. 4, Supplementary Table S2), with further speed-up over multiple cores, and is thus suitable for genome-scale analyses.

**Figure 4:**
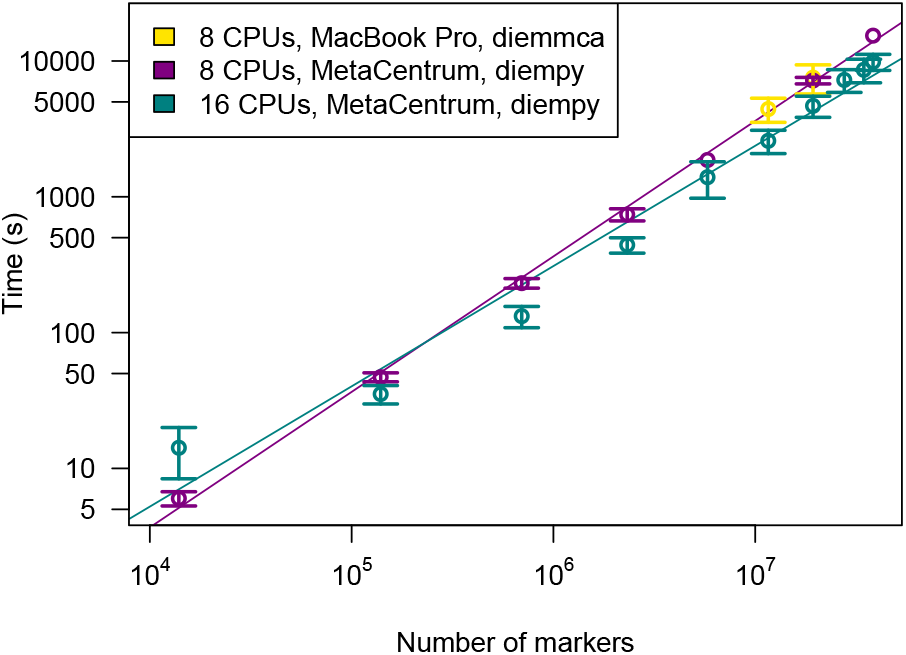
The *diem* computation time scales linearly with number of markers, and is reduced by making more CPUs available. Time trials were run on nested thinnings of the full mouse marker set, with replicates at each thinning level. Points and whiskers show mean ± SD in the thinning replicates. Yellow – absolute time of diemmca run on an 8 CPU Macbook Pro, purple (8 CPUs) and green (16 CPUs) – PBS monitor ‘job time’ for diempy jobs run on MetaCentrum computational cluster nodes (see Supplemental Table S2 for details), purple line – linear fit to all 8 CPU datapoints, rrespective of hardware, green line – linear fit to all 16 CPU datapoints, irrespective of hardware. Gradients 1.0728 × 10^*−*7^ and 7.0277 × 10^*−*8^ hrs/marker, respectively. Doubling CPU number gives ∼ 1.5 fold speed-up. diemmca and diempy results are identical on identical input. The Python version (diempy) currently requires more RAM resources due to differences in memory management between Mathematica kernels and Python parallel processes.

### Comparison with existing methods

Figure 5 overlays the introgressed blocks identified by Banker *et al*. (2022) onto the *diem* result in the *Vkorc1* region in individuals included in both studies. The major axis of difference between the studies is censorship of sites inspected. We suggest this is the effect mentioned in our Introduction: a bias against detection of geneflow results from declaring a priori that reference sets of individuals are ‘pure’ (Gompert *et al*. 2017). Banker *et al*. (2022) only inspect introgression into *spretus* at sites where selected individuals a priori labelled *domesticus* have invariant state; conversely, they only inspect introgression into *domesticus* at sites where individuals a priori labelled *spretus* have an invariant state. Therefore, all sites with intra-taxon variation are censored; All sites with bi-directional introgression are censored. From Figure 5 it appears this censorship leaves very few sites with which to measure sequence divergence. In contrast, *diem* has zero prior on the membership of genomes in taxa. Sites invariant across all genomes are censored, but this leaves large absolute numbers of variable sites (genome size × divergence) that can be inspected for introgression with an approach that is scalable over *n* markers (variable sites).

The comparison of the *diem* results to Structure shows that the posterior allele assignment to *domesticus*/*spretus* statistically matches the *diem* result (G-test: *G* = 2646211, χ^2^ *df* = 6, p < 0.001; Fig. 6). The major axis of difference between *diem* and Structure is that Structure imputes state whereas *diem* only polarises it. We compare Structure to *diem* using a contingency table summarising polarised vs imputed state for an interval of mouse chromosome 7 (Table 1). We see Structure imputation is consistent with *diem* homozygous states, but Structure overwrites unknown input states and tends to re-assign input heterozygous (1) state as homozygous (2). We can mimic this last aspect of the linkage model Structure imputation by re-calling all input states according only to the products of marginal genotype probabilities of individual genotypes and markers from genotypes polarised by *diem* (Table 1, *diem*-imputed). We see that imputation from *diem* marginals reproduces the tendency to re-assign heterozygous (1) input state as homozygous (2) in this dataset. We observe that the major difference between the *diem* marginals-only imputation and Structure linkage model imputation is a relative excess of Structure heterozygous calls. This is most noticeable for the individual with accession SAMEA3678857, which has majority input state homozygous *domesticus* (Fig. 3B), but after Structure imputation, the majority state is heterozygous (Fig. 6).

**Table 1:** Comparison of *diem*-polarized genotypes of mice with Structure SITEBYSITE linkage model genotypes in the partial chromosome 7 (cf. Fig. 3).

| *diem* polarisation | STRUCTURE |  |  | *diem*-imputed |  |  |
| --- | --- | --- | --- | --- | --- | --- |
|  | *domesticus* (0) | heterozygous (1) | *spretus* (2) | *domesticus* (0) | heterozygous (1) | *spretus* (2) |
| Undetermined (U) | 3,056,099 | 132,580 | 2,728,672 | 3,228,406 | 35,881 | 2,653,064 |
| *domesticus* (0) | 950,357 | 5,231 | 33,551 | 954,643 | 1,607 | 32,889 |
| heterozygous (1) | 30,370 | 55,433 | 122,237 | 76,070 | 24,048 | 107,922 |
| *spretus* (2) | 8,816 | 1,727 | 975,734 | 12,721 | 1,386 | 972,170 |

**Figure 5:**
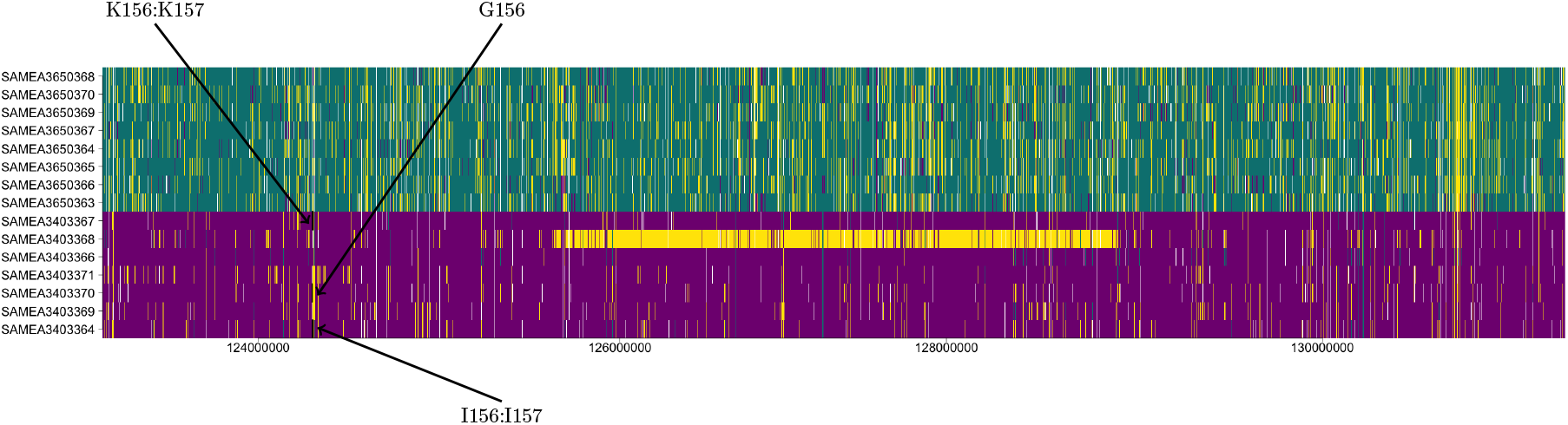
Comparison of *diem* genome polarization with a sequence-divergence based method to detect ntrogression (Banker *et al*. 2022). Partial mouse chromosome 7 genome polarisation corresponds to that in Fig. 3B, selecting individuals that were analysed both here and in Banker *et al*. (2022). Arrows point to black boxes delimiting introgressed regions identified in the respective cells of Suppementary Table 3 in Banker *et al*. (2022). The large, yellow block in sample SAMEA3403368 matches the ntrogression identified by Song *et al*. (2011) using Sanger sequencing.

**Figure 6:**
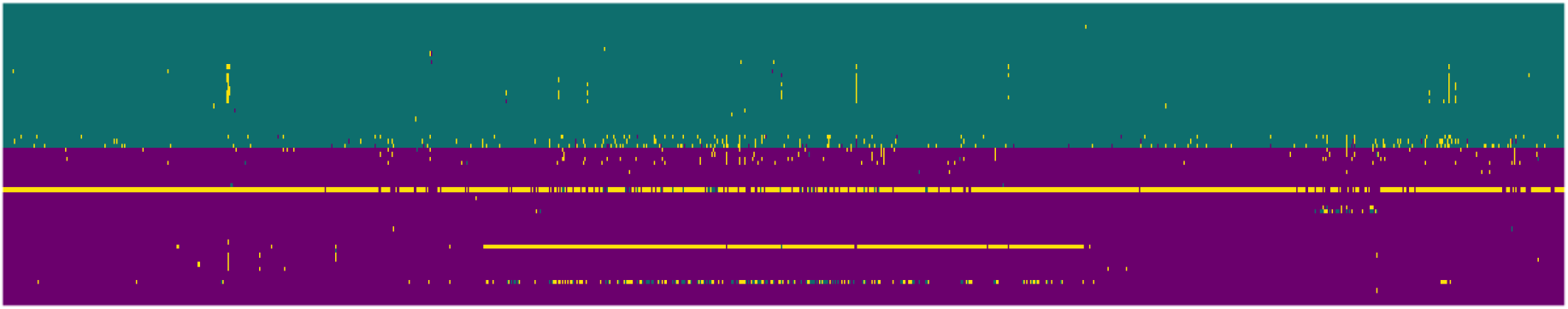
Genotype calls of mice from Structure SITEBYSITE linkage model. The individual ordering (rows) and colour encoding is identical to those in Fig. 3B. The horizontal, predominantly yellow line is an individual with accession number SAMEA3678857 that has been imputed as heterozygous by Structure (the transcriptomic data is very sparse in Fig. 3).

The mouse introgressions highlighted across the genome by *diem* are bidirectional (Fig. 7, Supplementary Table S3). While *M. musculus domesticus* gained a large introgressed block on chromosome 7 carrying resistance to warfarin, introgressions to *M. spretus* genome include genomic blocks on multiple chromosomes with genes involved in chemical sensing.

**Figure 7:**
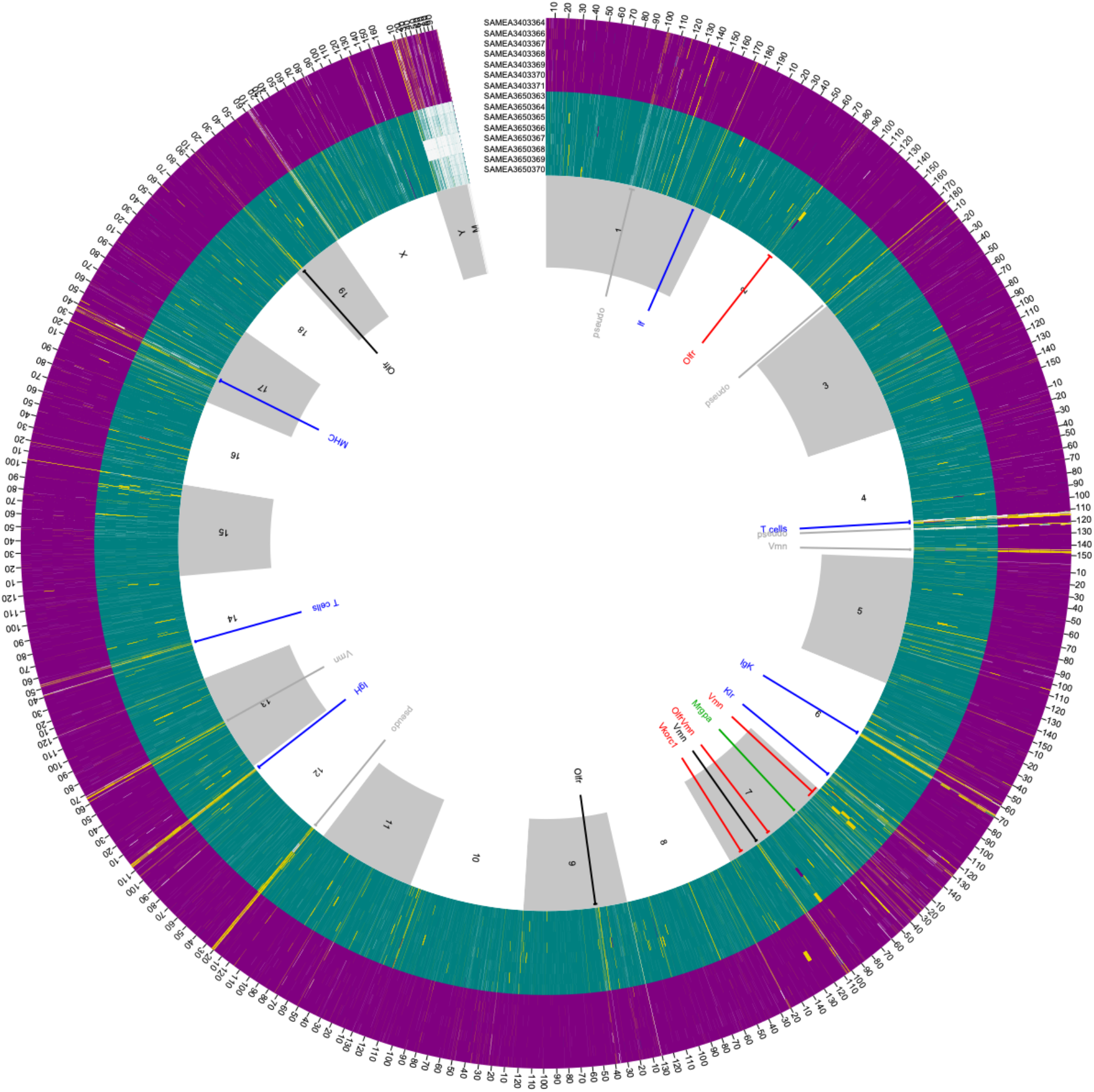
Visualising a mouse genome barrier: 38 million variable sites aligned to the GRCm39 reference and coloured using *diem* output. Purple – *M. musculus domesticus*, green – *M. spretus*, yellow – heterozygous or overmerged onto the reference, white – no call. Of the full *diem* analysis, only the fifteen ENA whole-genome submissions (accession numbers top, centre) are shown, ordered by their *diem*-estimated hybrid index. The transition from purple to green genomes is sharp, suggesting the genome barrier is strong. Inner grey/white ring – reference chromosome, outer ticks – reference chromosome position in Mb. Central annotation, major features: Red – regions of introgression ncluding *Vkorc1* ; the *M. spretus* version of this gene provides pesticide resistance to *M. musculus domesticus* carriers: SAMEA3403368 is a heterozygous carrier, *Vkorc1* being embedded in a ∼ 3.4Mb ong yellow block on purple background, demonstrating that the genome barrier is permeable. All other major introgressions are from *M. musculus domesticus* to *M. spretus* and occur in chemosensory gene clusters (Vmn: Vomeronasal, Olfr: Olfactory). Remaining annotated features: yellow – false heterozygous state due to overmerging, blue – wild mouse variation in immune gene clusters, and chemosensory gene clusters overmerges onto the GRCm39 reference, assembled with long reads from the lab mouse strain C57BL/6J, inbred for many hundreds of generations. Overmerged regions are associated with high pseudogenic C57BL/6J annotation density, sometimes to the extent that entire gene clusters are pseudogenic (grey). Supplementary Table S3 details the annotated regions.

## Discussion

Genome polarisation dissolves the concept of populations: they exist nowhere in the *diem* algorithm. This might seem surprising for a method inspired by population genetics, but allowing for continuous change in genotypes in fact dates back to the neo-Darwinian synthesis (Haldane 1948), while inferring barriers to geneflow from abrupt change in correlated genotypes (Barton 1979b,a) lies at the base of (arguably) the most replicable and productive treatment of biological field data: hybrid zone analysis. We have shown that the *diem* algorithm is robust in detecting a single homogeneous set of individuals (Fig. 2) and a barrier between two taxa (Fig. 3), including detection of introgressed blocks (Figs. 3B, 7), *diem* allows analyses of samples with mixed levels of reduced genome representation, the implementation scales to genome-size data (Fig. 4), and diemr, diempy and diemmca provide matching solutions that make the approach available to a wide potential userbase.

Polarising genomes allows us to study a continuous genomic space, investigating barriers to geneflow and the capture of genetic material at a regional scale. An emphasis on sequences (along-genome summaries) in previous phylogenetic-inspired approaches is replaced by an emphasis on markers (across-genome summaries). Perhaps most promisingly, *diem* informs hypothesis formation at the genome scale. For example, we can formulate a new hypothesis for the across-mouse-species admixture event analysed here. An event seen as unidirectional capture of pesticide genes with some evidence of bidirectional geneflow can be reconsidered as an exchange of genes suited to the anthropocene. While long-time synanthropic *M. musculus domesticus* received pesticide resistance from *M. spretus*, which previously lived further from human influence, no species can now live far from humans, and *M. spretus* may benefit from sets of chemosensory genes honed by *M. musculus domesticus* in human proximity.

### Designing *diem* studies

For those familiar with classic hybrid zone studies, and having sufficient resequencing budget, *diem* simplifies classic design. The preliminary step of defining reference sets and surveying potential markers for diagnosticity between those sets is dropped. Individuals from the hybrid zone are resequenced, the data is aligned, and polymorphic sites polarised. The zone is then characterized using polarised markers of high diagnostic index. Cline inference can be checked for invariance over definitions of ‘high’. A more complete replication of the classical design constructs reference sets, but chosen post-*diem*, using individuals’ estimated hybrid indices. The zone is then, as in the classical case, characterized using markers that diagnose these reference sets, polarised with respect to them. In this more complete replication, we see choice of reference sets is now informed by pre-knowledge of their genome content rather than ad hoc based, for example, on sampling locations in the field. The first design is further applicable even if field sampling discovers no pure individuals. Both may recover many more sites of the genome than classical for downstream analyses of clinal patterns. For those analyses, *diem* provides the copolarisation of genome state encoding across loci implied by ‘similar’ in paragraph two of Barton (1979b): *“Suppose there is a cline at some locus, maintained at equilibrium by natural selection in the face of dispersal; alleles at any locus closely linked to this cline, and themselves more or less neutral, will tend to show a similar pattern of variation”* and ‘the same’ in section ii (c) of Barton (1979a): *“Suppose that a hybrid zone is maintained by epistatic selection between two loci [*…*] The two clines will have the same shape (by symmetry)*.*”* Further, while here we describe genome wide correlations generated by divergence, we could equivalently talk in classical terms of the genome wide linkage disequilibrium generated by admixture, either natural or artificial, *sensu* Baird (2015).

Potential applications of *diem* are much wider. First, regarding input data: While these classical scenarios are reduced genome representation approaches, the reduction step is post *diem*, and so a large resequencing budget may be incurred. Reduced representation genomics is often equated with budget reduction, and *diem* applies not just to resequencing data, but also to approaches which subsample the genome such as RNAseq, RADseq, SNP chips and mixed data from any and all such sources (as with our mouse example). *diem* also naturally extends to long read data. As with short read data, within-read phasing is ignored. All data types are aligned and polymorphic sites identified. Encoding polymorphic sites from mixed data is straightforward because missing data is part of the *diem* input format and imparts no bias. Because polarisation signal is associative, a general rule of thumb is: the more *diem* input genomes, the better, irrespective of format mixing. Users should however be cautious of SNP chip data. Some SNP chips are designed not only to target potentially polymorphic sites, but sites assumed to diagnose pre-defined reference sets, risking re-introducing the bias against detection of introgression cautioned by Gompert *et al*. (2017).

Second, regarding design hypotheses: Because no prior information regarding barriers, taxonomy or genome architecture is needed, *diem* is suited to analyses of non-model, poorly characterized and cryptic systems. When a barrier analysis includes a cryptic outgroup genome in error, its presence is evident when markers are ordered along genomes as with Fig. 7: The barrier divergence history is not part of the outgroup’s divergence history, and so when outgroup genetic states are polarised with respect to the barrier they assign to barrier side at random. In large *n* plots such as that in Fig 7, they then appear as 4-colour blends centred on h *∼* 0.5.

The lack of required priors for *diem* extends to ignorance of ploidy. First, the *diem* state encoding generalises without alteration to ploidy-free definitions ‘one state present’ for the ‘homozygous’ (0,2) encodings and ‘both (commonest) states present’ for the ‘heterozygous’ (1) case. Higher ploidies will simply tend to increase the (U,1) state frequencies, reducing signal without biasing polarization. Second, if the researcher supplies a uniform compartmental ploidy to *diem*, its absolute ploidy level cancels at its only point of use: the calculation of hybrid indices (Eq. 7). The possibility of inputting heterogeneous ploidies exists only to allow that, for example, the single X of a male contributes only half as much to its hybrid index estimate as the two Xs of a female. Under this slight approximation, preliminary analyses with a uniform ploidy prior may even help identify sex chromosomes.

Consider *diem* output resulting in a visualization resembling that of the bat dataset here, except for one genome region that instead resembles that of the mouse dataset. This would indicate a genome-localized barrier, perhaps reflecting reduced recombination in an inversion or close to a centromere, perhaps increased barrier strength at a magic gene or, if the region encompasses a chromosome, increased barrier strength between sex chromosomes due to Haldane’s rule. This wide potential for *diem* applications is attractive but requires some caution. While interpretation of the classical examples above may be regarded as well validated because *diem* merely provides reduced-bias data to a well understood inference scheme in use for three decades, interpretation of genome-localised barrier signal is less well developed. In such cases, *diem* users may wish to simulate heterogeneous genome scale divergence for hypothetical scenarios, and compare the polarised simulation output to signal in their own data as a validation step. Further caution is required here. There is no strong correspondence between simulating K barriers (strength measured in units distance) and simulating K + 1 population genetic clusters. As is made clear in the Structure manual (section 5.4) *“The underlying structure model is not well suited to data from this [isolation by distance (IBD)] scenario”* (Pritchard *et al*. 2010). Simple examples disprove the correspondence. A ring species is a single IBD population, yet possesses a barrier amenable to *diem* analysis. A hybrid zone crossed by two geographic barriers is amenable to *diem* analysis with two ‘pure colours’ (the stronger barrier will be taken as ‘central’). While posthoc comparison of Structure analyses may indicate three clusters, *diem* will show a single cline with two steps.

While this lack of correspondence between clusters and barriers may force a shift of study design away from cluster- and toward barrier-based-thinking, this may be positive. In his perspective on species definitions for the modern synthesis (Mallet 1995) summarises Wallace’s critique: *“Wallace presented a very clear interbreeding species definition, then immediately dismissed it [*…*]. ‘Species are merely those strongly marked races or local forms which, when in contact, do not intermix, and when inhabiting distinct areas are*…*incapable of producing a fertile hybrid offspring. But as the test of hybridity cannot be applied in one case in ten thousand, and even if it could be applied, would prove nothing, since it is founded on an assumption of the very question to be decided*…*it will be evident that we have no means whatever of distinguishing so-called “true species” from the several modes of [subspecific] variation here pointed out, and into which they so often pass by an insensible gradation*.*’”* Mallet (1995) continues, *“Wallace is first saying that it is practically impossible to make all the necessary crosses to test genetic compatibility. Second, since theories of speciation involve a reduction in ability or tendency to interbreed, species cannot themselves be defined by interbreeding without confusing cause and effect*.*”* We would add first, even if we cannot test all cases for barriers, this does not reduce the utility of any one such test, nor does the comparative method require all possible comparisons, and second, detecting a barrier separating genomes in the absence of any prior taxonomic information is *not* founded on an assumption of the very question to be decided.

## Acknowledgements

We thank Beate Nürnberger for discussions and constructive criticism. SJEB thanks the Lohse Lab, University of Edinburgh, Scotland, and the Barton Lab, IST, Austria, for discussions during development. We would like to thank Nick Barton for comments received during revision of the manuscript. In 1996 and 1999, SJEB worked in Montpellier side by side with Annie Orth, who deserves credit for the initial discovery of geneflow between *M. musculus domesticus* and *M. spretus*.

This work was supported by the Institute of Vertebrate Biology of the Czech Academy of Sciences (RVO: 68081766), and Czech Science Foundation “We Support Fundamental Research”, RECETOX Research Infrastructure (No. LM2018121) financed by the Ministry of Education, Youth and Sports, and the Operational Programme Research, Development and Education (the CETOCOEN EXCELLENCE project No. CZ.02.1.01/0.0/0.0/17 043/0009632). MetaCentrum computational resources were supplied by the project “e-Infrastruktura CZ” (e-INFRA CZ LM2018140) supported by the Ministry of Education, Youth and Sports of the Czech Republic.

## Author Contributions

Conceptualization: SJEB; Methodology: SJEB, NM; Software: SJEB, NM, JP, IJ; Validation: SJEB, NM, PŠ; Writing - original draft: SJEB, NM; Writing - review & editing: SJEB, NM, JP, IJ, PŠ.

## Data Availability

The data underlying this article are available in the GenBank SRA Database at https://www.ncbi.nlm.nih.gov, and can be accessed with BioProject and BioSamples identifiers listed in text and in the Supplementary Table S1.

## Code Availability

The *diem* method is implemented in Mathematica, Python and R. The Mathematica and Python codes are available at github through https://github.com/StuartJEBaird as diemmca and diempy, respectively, and the R package diemr is available at CRAN through https://CRAN.R-project.org/package=diemr.

## Supplementary Material

Table S1: Tab-separated table of BioSample Accession Numbers, geographic origin and hybrid indices (HI) of *Mus musculus domesticus* and *Mus spretus* analysed by *diem*.

Table S2:Comma-separated table with benchmarking results of *diem* implemented in diempy on the MetaCentrum computational cluster.

Table S3:Tab-separated table with genomic regions introgressed and overmerged between *M. muscuus domesticus* and *M. spretus* shown in Fig. 7. Positions refer to mouse genome reference GRCm39, NCBI Assembly Accession: GCA_000001635.9.

